# Bacterial genome architecture shapes global transcriptional regulation by DNA supercoiling

**DOI:** 10.1101/561423

**Authors:** Bilal El Houdaigui, Raphaël Forquet, Thomas Hindré, Dominique Schneider, William Nasser, Sylvie Reverchon, Sam Meyer

**Affiliations:** Université de Lyon, INSA Lyon, Université Claude Bernard Lyon 1, CNRS UMR5240, Laboratoire de Microbiologie, Adaptation et Pathogénie, 69621 Villeurbanne, France; Université Grenoble-Alpes, CNRS UMR5525, TIMC-IMAG, 38000 Grenoble, France

## Abstract

DNA supercoiling acts as a global transcriptional regulator in bacteria, that plays an important role in adapting their expression programme to environmental changes, but for which no quantitative or even qualitative regulatory model is available. Here, we focus on spatial supercoiling heterogeneities caused by the transcription process itself, which strongly contribute to this regulation mode. We propose a new mechanistic modeling of the transcription-supercoiling dynamical coupling along a genome, which allows simulating and quantitatively reproducing *in vitro* and *in vivo* transcription assays, and highlights the role of genes’ local orientation in their supercoiling sensitivity. Consistently with predictions, we show that chromosomal relaxation artificially induced by gyrase inhibitors selectively activates convergent genes in several enterobacteria, while conversely, an increase in DNA supercoiling naturally selected in a long-term evolution experiment with *Escherichia coli* favours divergent genes. Simulations show that these global expression responses to changes in DNA supercoiling result from fundamental mechanical constraints imposed by transcription, independently from more specific regulation of each promoter. These constraints underpin a significant and predictable contribution to the complex rules by which bacteria use DNA supercoiling as a global but fine-tuned transcriptional regulator.

## 1 Introduction

The role of DNA supercoiling (SC) in transcriptional regulation has attracted considerable attention in recent years. Due to the helical nature of DNA, mechanical torsion affects transcription at both initiation and elongation steps, and can thereby be considered as a non-conventional transcriptional regulator in eukaryotes as well as bacteria [1, 2, 3, 4]. In the latter, fast changes in DNA topology play a central role in the global transcriptional response to environmental stress [3, 5]. Inheritable changes in DNA topology are also under positive selection during evolution experiments with bacteria, in which SC-modifying mutations can provide a substantial fitness gain [6]. The regulatory action of SC is usually analysed from transcriptomes obtained after treatment by DNA gyrase inhibitors, causing global relaxation of the chromosome and changes in the transcription level of hundreds of genes [7, 8, 9, 10]. Since topoisomerases are found in all bacterial species, including those almost devoid of transcription factors such as *Mycoplasma* or *Buchnera* [11, 10], they might underpin an ancestral and widespread global regulation mode. However, our understanding of the underlying mechanisms remains limited, as there is currently no quantitative or even qualitative systematic model of the features defining a supercoiling-sensitive gene, and how it responds to SC changes. Here, we hypothesise that the local genomic architecture (*i.e.*, relative orientations and distances between adjacent genes) is a primary factor in this response, by dictating the distribution of SC at the gene’s promoter. Indeed, as recognised more than 30 years ago [12], transcription also generates significant torsional stress during the elongation step [13]. This stress is positive in front of the elongating RNA polymerase (RNAP) and negative behind it, and is able to diffuse along the doublehelix at distances of several kilobasepairs and reach nearby promoters [12, 14]. The relationship between transcription and SC is thus double-sided, and constitutes a dynamic and spatially organised coupling [15], hereafter quoted Transcription-Supercoiling Coupling (TSC). TSC has been proposed to underpin a complex and nonlinear interaction between adjacent genes depending on their relative orientations [15, 16, 17, 18], which could play a fundamental yet unexplored role in the supercoiling regulation mode of gene expression [5].

Several TSC models have been recently proposed [15, 19, 16, 17] in a biophysical and essentially theoretical perspective, *e.g.*, aiming at reproducing so-called “transcription bursts” [20]. But strikingly, no attempt has been made so far to simulate any specific experimental system, for which these models lack important components (*e.g.*, explicit topoisomerase enzymes). The new TSC model presented here is inspired by previous ones in that it describes the 1D stochastic binding and elongation of RNAPs along genes and the SC distribution resulting thereof; however, in contrast to these previous models, it was specifically developed and applied in a regulatory perspective, *i.e.*, with the aim of simulating actual biological systems and quantifying the effect of TSC-mediated regulation *in vivo*. It includes new realistic modeling ingredients that allow (i) precisely mimicking specific experimental assays through a small number of biologically relevant input parameters (promoter initiation rates, topoisomerase concentrations, position of topological barriers), (ii) directly comparing the results of the simulations to observations, and (iii) inferring the underlying dynamics of transcription and propagation of supercoils. In this paper, we use this model to develop the first systematic comparison between TSC simulations and gene expression data.

After presenting the model, we first simulate a series of *in vitro* or *in vivo* experimental systems on SCsensitive model promoters. We show that our simplified description is able to reproduce the quantitative effect of TSC on gene expression on the chromosome, and demonstrate that it is largely dictated by local gene orientations. We then propose that the genomic context may be a strong determinant of the “supercoiling-sensitivity” of many bacterial genes, independently from any sequence specificity of their promoter. We analyse existing and new transcriptomic data obtained in conditions of gyrase inhibition by antibiotics causing chromosomal relaxation, and show that convergent genes are significantly more activated than divergent ones in several bacterial species. We then demonstrate that this behaviour results from the basic mechanical constraints imposed by transcription, independently from species- or gene-specific properties. These constraints define how DNA topology, globally controlled by the cell physiology, affects the expression of genes according to their local orientation, promoter strength and distance. Finally, we ask if this form of genome-printed regulation can contribute to bacterial evolvability; we analyse global transcription profiles obtained from the longest-running evolution experiment, in which SC-modifying modifications have been selected. As predicted by our TSC modeling, we demonstrate that genes’ expression changes in the evolved strains with modified SC are related to their local orientation. This analysis suggests that the regulatory rules dictated by neighbour genes’ topological interactions likely constitute a robust and fundamental constraint governing the evolution and regulation of bacterial genomes.

## 2 MATERIALS AND METHODS

### 2.1 Model equations

Our model describes the dynamic transcription-supercoiling coupling. Most hypotheses and components of the model are described in Results and Discussion; here, we provide equations and parameter values. The promoter response curve (Fig. 1C) is computed from a thermodynamic model of transcription, *f* (*σ*) = exp(*m U* (*σ*)) where *U* (*σ*) is the SC-dependent promoter opening free energy. It follows a sigmoidal curve [15]: *U* (*σ*) = 1*/*(1 + exp((*σ* -*σ*_*t*_)*/ϵ*)). *σ*_*t*_ is the threshold of promoter opening, *ϵ* sets the width of the crossover, and 1*/m* is an effective thermal energy that sets the SC factor. Standard values shown on Fig. 1C are *σ*_*t*_ = −0.042, *ϵ* = 0.005, *m* = 2.5 (calibrated on the *pelE* promoter, see below).

**Figure 1:**
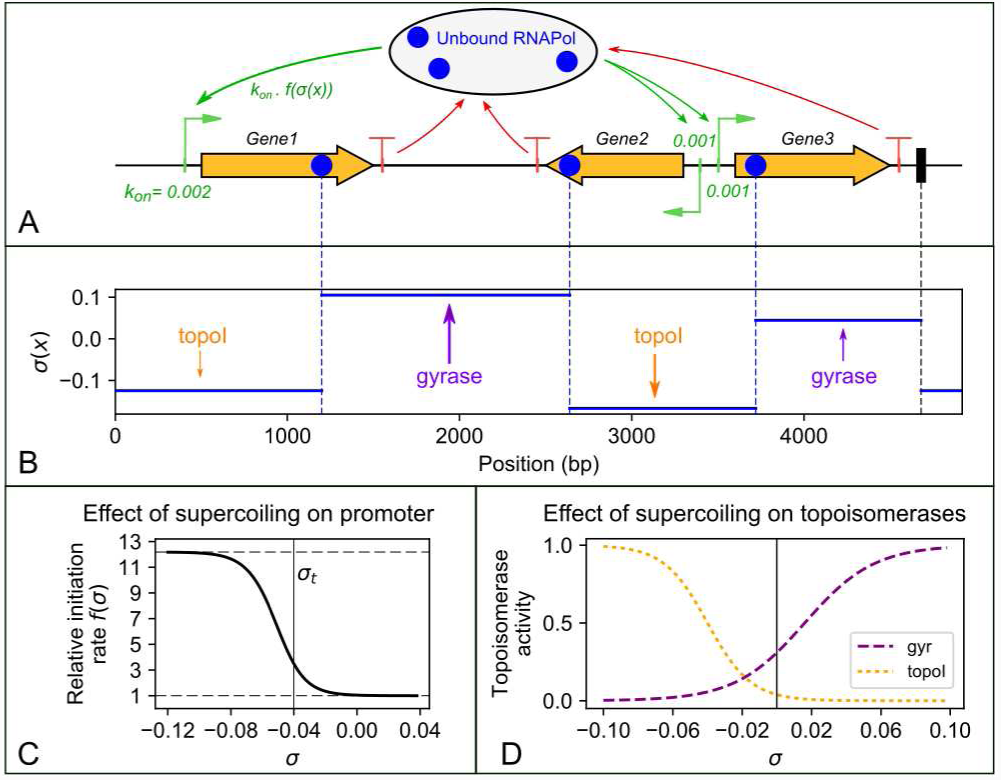
Illustration and main components of the transcription-supercoiling coupling model. **(A)** Snapshot of the simulation of the stochastic binding (green arrows; the basal initiation rate *k*_*on*_ of each promoter is shown), elongation, and dissociation (red arrows) of a set of RNAPs along a 1D genome (here a 5kb plasmid). **(B)** The SC profile is updated at each timestep, and is affected by elongating RNAPs as well as by topoisomerase activity. This level is constant between topological barriers, *i.e*., either elongating RNAPs (blue) or fixed proteic barriers (black). **(C)** The local SC level affects each promoter through an activation curve derived from thermodynamics of open complex formation, which modulates its specific strength (basal initiation rate). **(D)** Topoisomerases bind in a deterministic but heterogeneous way, according to the local SC level (see text).

Topoisomerase activity curves *t*(*σ*) (Fig. 1D) follow sigmoidal curves (same equation as above) parameterised from experimental assays [21, 22]: bacterial topoisomerase I acts on negatively SC DNA only, whereas the DNA gyrase exhibits moderate (∼ 30%) activity on relaxed DNA, and is fully active at σ ≃ 0.1. Accordingly, thresholds of the sigmoids were set at values −0.04 and 0.01, and crossover widths at 0.012 and 0.025 respectively (Fig. 1D). Basal activity constants were calibrated on *in vitro* assays of transcription-induced SC accumulation (Fig. 2B, see Results): *k*_*topo*_ = ±0.001*s*^-1^ (for topoisomerase I and gyrase respectively).

**Figure 2:**
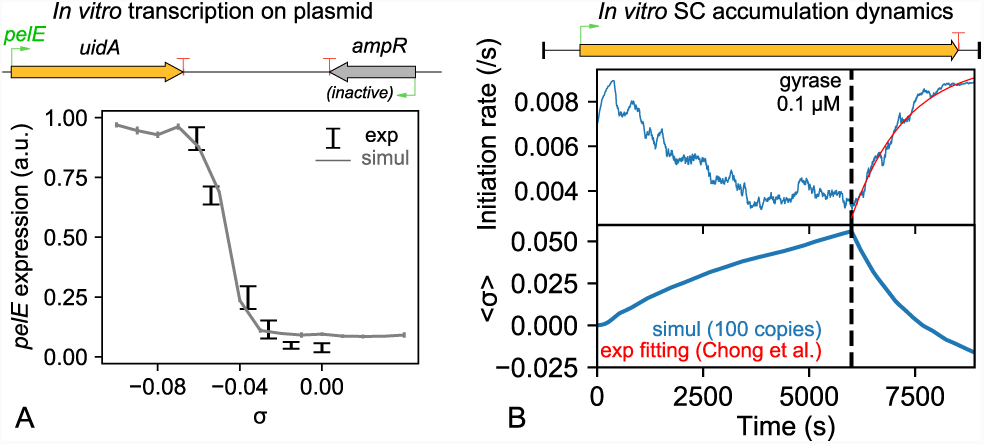
Calibration of the model on *in vitro* transcription experiments with plasmids. **(A)** The promoter activation curve (Fig. 1C) is calibrated from *pelE* expression levels measured on purified plasmids prepared at different SC levels [3]. Due to the absence of topological barriers in the plasmid, transcription-induced supercoils do not accumulate and SC levels remain constant. In this assay, this promoter from *Dickeya dadantii* is activated around 20-fold by negative SC without any addition of transcription factors. **(B)** Topoisomerase activity constants are calibrated from a SC accumulation assay [20]. The instantaneous initiation rate (top) was measured over time, and is reproduced by our simulations where the average plasmid SC level can be inferred (bottom). Positive supercoils first accumulate in the absence of DNA gyrase, resulting in a progressive repression of the promoter, and are then released in an exponential timecurve (red curve) reflecting gyrase activity.

### 2.2 Simulation methods

The genome was discretised with a 60-nt unit length, and simulations were performed with the Euler algorithm using a constant timestep *dt* = 2*s*. At each timestep, free RNAPs are stochastically assigned to the available promoters according to their instantaneous initiation rate, or stay unbound; elongating RNAPs are moved by one unit length (speed 30 nt/s), and unbind when they reach a terminator. The SC levels of all topological regions (separated by fixed proteins or RNAPs) are then updated. Each elongating RNAP adds a constant number ±Γ of supercoils (in the regions ahead or behind it, respectively), which is normalised by the region length to compute the increment in superhelical density. Γ represents the fraction of DNA coils that are converted into supercoils by the elongating RNAP, and might depend on the environment (molecular crowding, viscosity); a value of 0.2 / unit length was used in all simulations presented here. Topoisomerases then increment the SC level of each domain by *k*_*topo*_ *t*(*σ*) *dt*. In the absence of topoisomerases, we checked that the genome-averaged SC level is constant throughout the simulation. Simulations, as well as all data analyses, were carried in Python with the NumPy numerical package. Note that the effective initiation rate of a gene in a simulation depends not only on its basal rate, but also on the number of RNAPs in the simulation, SC levels, and initiation rates of the other promoters. In all simulations of *in vivo* systems, we used a concentration of 0.25 *µ*M for gyrase, and 0.025 *µ*M for topoisomerase I [20].

In our description, nucleoid-associated proteins can act only as topological barriers or by modulating promoter strengths (*i.e.*, in the role of global transcription factors), while their effect on SC within topological domains is neglected; since a notable part of this effect consists in modifying the twist/writhe equilibrium of SC, this hypothesis is consistent with our unidimensional description of DNA [2, 23] (see below). Similarly, we disregard the presence of specific sequences favouring structural transitions of DNA [24, 25] or DNA gyrase binding.

### 2.3 Transcriptomics data

Transcriptomes of *Dickeya dadantii* were analysed as previously described [9]. *D. dadantii* 3937 cells were cultivated in M63 minimal medium supplemented with 0.2% (wt/vol) sucrose as carbon source, with or without 0.2% (wt/vol) polygalacturonate (PGA, a pectin derivative). Cells were harvested in early exponential phase (OD_600_=0.2, sucrose) or transition to stationary phase (OD_600_=1.5 in sucrose and OD_600_=1.8 in sucrose+PGA). For each condition, total RNAs were extracted using the frozen-phenol procedure [26], either from untreated cells or from cells treated with 100 *µ*g/mL novobiocin for 15 min, with two biological replicates. At this concentration, novobiocin has no impact on *D. dadantii* growth [3]. Control experiments for RNA extraction quality, absence of DNA contamination, and qRT-PCR validation of selected genes were conducted as previously [9]. Further steps were carried out by Vertis Biotechnologie AG (http://www.vertis-biotech.com): rRNA depletion using the Illumina Ribo-Zero kit, RNA fragmentation, strand-specific cDNA library preparation, and Illumina NextSeq500 paired-end sequencing (∼15 million paired reads per sample).

Transcriptomes of *Escherichia coli* clones isolated from the Long-term Evolution Experiment (LTEE) [27, 28] were obtained as previously described [29]. Briefly, bacterial strains were grown in LTEE conditions (Davis minimal medium containing 25 *µ*g/mL of glucose) but in 200 mL cultures to obtain sufficient amounts of RNA. Bacteria were harvested at mid-exponential phase and total RNAs were extracted from cells pellets using Qiagen’s RNeasy Cell Tissue Kit, following the manufacturer’s protocol. RNaseOUT (Invitrogen) was added to the RNA extracts that were subsequently handled and analysed by Beckman Coulter Genomics according to the standard Affymetrix GeneChip protocol for bacteria, and using the Affymetrix *E. coli* Genome 2.0 Microarray. Quality checks and normalisation were performed using the Affymetrix GeneChip Command Console and Expression Console according to standard Affymetrix protocols. The three strains used in this study were the ancestral strain of the LTEE, REL606, and two clones previously isolated from the Ara-1 population at 2,000 (REL1164A, quoted “2K” below) and 20,000 (REL8593A, quoted “20K” below) generations [30]. For each strain, 6 independent cultures were grown and 6 RNA extracts were analysed (18 microarrays in total).

### 2.4 Statistics and data analysis

Sequenced reads from *D. dadantii* were mapped on the reference genome (NCBI NC_014500.1) with Bowtie2 [31], counted with htseq-count [32] against the reference annotation, and gene differential expression was analysed with DESeq2 [33]. A threshold of 0.05 on the adjusted p-value was chosen to define differentially expressed genes (between 1250 and 1750 for novobiocin treatment).

The normalised dataset from *E. coli* (10,208 probes per microarray) was cured to only retain data from Affymetrix probes that entirely match within one coding sequence of the REL606 reference genome (NCBI NC_012967.1) and with the highest score of sequence homology in case several probes target the same gene. The cured dataset (3839 probes per assay) was obtained for all three strains (6 replicates each) and log2 transformed. Differential expression was analysed with the optimal discovery procedure [34] implemented in the EDGE v1.1.290 software [35], and based on the false-discovery rate. Differentially expressed genes were first defined using a q-value threshold of 0.1; for the orientation analysis which required larger datasets (see below), a looser threshold of 0.25 on the p-value was used instead.

All processed data are available in a Supplementary File. The relation between gene response and local orientation was carried with a homemade Python code. All error bars shown are 95% confidence intervals. Proportions were compared with the *χ*^2^ test. Note that when we discuss the chromosomal SC level, the expression “increased SC” refers to the absolute level (although it is here negative), as often in microbiological literature.

## 3 RESULTS AND DISCUSSION

### 3.1 Stochastic modeling of the Transcription-Supercoiling Dynamical Coupling

The transcription-supercoiling coupling (TSC) is complex and dynamic, and must therefore be simulated with a suitable biophysical model. As previously proposed [15, 16, 17], we simulated the transcription dynamics of a circular genome (which can be a plasmid, but also an entire chromosome) as a unidimensional system, as illustrated on Fig. 1 in a linearised depiction. At each timestep, (A) RNA Polymerases (RNAPs) bind stochastically at promoters (with different basal initiation rates *k*_*on*_), transcribe the genes, and eventually unbind at terminators; and (B) the local distribution of SC is affected by transcription and by the action of the two major topoisomerases, DNA gyrase and topoisomerase I. These two aspects of the genome (gene expression and physical state) affect each other in a reciprocal way: (i) elongating RNAPs act as topological motors that pump positive supercoils from back to front, resulting in heterogeneous SC distributions; (ii) the initiation rate of each promoter is modulated by the local level of SC, following a single response function (Fig. 1C). Qualitatively, these two effects are sufficient to generate a local form of transcriptional regulation mediated by SC [15, 16]. For a quantitative modeling of the process, a series of simplifications, hypotheses and additional components are required, as follows.

Within our unidimensional description, only the *twist* (torsional) contribution to SC is considered, rather than the complete (*twist + writhe*) level that involves 3D deformations of the double-helix [36]. This approximation is justified as torsional deformations are the main modulators of the free energy of DNA melting required for transcription initiation, which may constitute the major mechanism of regulation by SC *in vivo* [2, 15]. We assume that this regulation follows a single response curve (Fig. 1C), obtained from the thermodynamic opening curve of a bacterial promoter [3, 15]. The elongation speed is assumed to be constant (measured values are indeed relatively homogeneous [37]), and to give rise to a constant rate of SC generation. Note that the stalling effect of positive supercoils on the elongating RNAP is thereby neglected, which is likely relevant for most moderately transcribed genes in the globally underwound bacterial genome, although possibly inaccurate for very strong promoters like those of ribosomal RNAs [17]. As previously proposed [17, 23], RNAP-generated super-coils are assumed to diffuse instantaneously (at the timescale of the simulation, *i.e.*, seconds) along kilobase distances, until reaching another RNAP or a fixed topological barrier, which can represent H-NS or other DNA-bound proteins with the same ability to isolate supercoils [2, 5]. The resulting instantaneous SC profile (Fig. 1B) appears as a succession of flat regions, in contrast to the more continuous and heterogeneous profiles expected if the diffusion was slower [16]. Finally, the activities of topoisomerases are considered as deterministic and continuous, whereby DNA gyrase introduces negative supercoils and topoisomerase I relaxes DNA with nonlinear activation curves calibrated from experimental knowledge (see Fig. 1D and Materials and Methods), without any sequence preference. As a result of these simplifications and hypotheses, the proposed model involves only a small number of mechanistic parameters, all of which are either obtained from experiments or calibrated on *in vitro* transcription assays, as follows (equations in Materials and Methods).

The promoter response curve was calibrated from the *pelE* promoter (Fig. 2A), which encodes a major virulence factor of the phytopathogenic enterobacterium *Dickeya dadantii*, and is strongly SCsensitive [3]. In this assay, plasmids carrying a *pelE* promoter were prepared at different SC levels; since they were free to rotate in solution and carried no topological barrier, the supercoils generated by transcription merged instantaneously, and the SC level thus remained constant with time. The measured expression levels thus directly reflected the promoter response curve (Fig. 1C) and match the dependency expected from promoter opening thermodynamics [15]. Conversely, in order to calibrate the dynamics of SC accumulation resulting from topoisomerase activity, we simulated an experiment involving a plasmid with a long (12 kb) gene, anchored to a surface that hampers rotation (Fig. 2B and [20]). Topoisomerase I was first introduced at a well-controlled concentration (41 nM): as a result, during each elongation event, only the negative twin-domain was relaxed by topoisomerase I, whereas the positive supercoils were not relaxed; this unbalance resulted in a slow accumulation of positive supercoils (in a timescale of hours), and promoter repression (by a factor 3). The sudden addition of DNA gyrase (at 0.1 *µ*M) then restored the initiation rate within less than one hour. After calibrating the topoisomerase activity constants, simulations closely reproduce the observed behaviour, and allow inferring the underlying SC level of the plasmid. Note that in this artificial construct involving strongly positive SC levels, the latter probably hampers transcription not only at the initiation step, but also during elongation [20]; accordingly, the standard initiation curve (Fig. 1C) had to be replaced by an effective curve repressed at positive SC levels (Fig. S1). We now show that in spite of its simplicity, our model is able to reproduce quantitative measurements of TSC-induced transcriptional interaction between adjacent genes in well-controlled *in vivo* experimental setups.

### 3.2 Torsional interaction between adjacent genes *in vivo*

The asymmetric nature of TSC implies that adjacent genes interact in a strongly orientation-dependent manner: divergent genes are expected to activate each other, whereas convergent genes experience a mutually repressive interaction [15]. To test this effect, Sobetzko [13] inserted a cassette containing two closely located divergent genes into the *Escherichia coli* chromosome (Fig. 3A), one of them being inducible (*tetP*). When the inducer concentration was varied, the expression of both genes changed in a coordinated manner, which was attributed to an activation of the second (non-inducible) gene by SC. We simulated this construct, using its exact architecture and physiologically relevant parameters (see Materials and Methods). Simulations reproduced a mutual activation when the basal level of *tetP* was progressively increased (Fig. 3A, right panel). Interestingly, since both promoters are always located in the same topological domain and experience the same SC level in our simulations, the observed increase in the expression of *tetP* results not only from the inducer concentration, but also from the resulting increase in negative SC in the central region (that is responsible for *tufB* activation). As can be observed, the activation of *tetP* is only semi-quantitatively reproduced, as the simulated levels remain below experimental ones: this small discrepancy may be indicative of a different SC-sensitivity for the two promoters (weaker for *tufB*), in contrast to our unique response curve (Fig. 1C).

**Figure 3:**
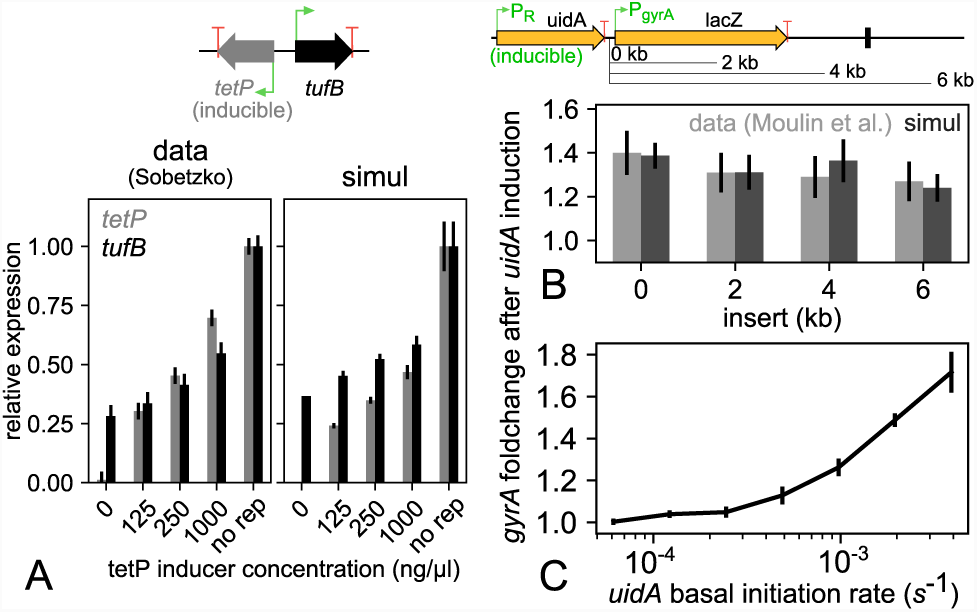
*In vivo* TSC-induced co-regulation of adjacent genes on the *Escherichia coli* chromosome. **(A)** The expression of the inducible *tetP* promoter (by increasing *tet* inducer concentration or absence of *tet* repressor) progressively activates a divergent promoter (*tufB*) [13]. This effect is reproduced semiquantitatively in our simulations by only changing the *tetP* basal initiation rate; small discrepancies with the data may reflect a different SC-sensitivity of the two promoters (see text). **(B)** The *gyrA* promoter is activated by the expression of the upstream isodirectional gene *uidA*, almost independently of the distance between the two genes (from 0 to 6 kb) [14]. This observation is fully consistent with our hypothesis of fast diffusion of SC over kilobase distances. A reversed promoter activation curve was used here to account for the unusual properties of the *gyrA* promoter (see text and Fig. S2). **(C)** Simulations (without insert) show that the *gyrA* activation factor is strongly dependent on the expression strength of the *uidA* gene.

This experiment shows that 300-bp distant promoters can significantly influence each other through TSC; however, can the resulting supercoils then diffuse at kilobase distances, as relevant to most genomic loci? Moulin et *al*. [14] measured the induction of the *gyrA* promoter by an upstream inducible gene located at various distances up to more than 6 kilobases (Fig. 3B). Note that *gyrA* is a very unusual promoter, as its SC-sensitivity goes opposite to most other promoters with an activation by DNA relaxation, which ensures homeostasis of the SC level in the cell. In the considered construct, this activation is produced by the positive supercoils resulting from the activity of the upstream *uidA* gene. Strikingly, the activation level is almost independent of the distance between the two genes, showing that TSC may be able to couple most genes located within topological domains of 10-20 kilobases [38], probably resulting in complex collective interactions. By construction (instantaneous diffusion), our model allows SC to diffuse at kilobase distances until it reaches a topological barrier; it is therefore no surprise that the experimental profile is easily reproduced under physiological topoisomerase concentrations (Fig. 3B), using an unusually reversed promoter activation curve relevant to *gyrA* (Fig. S2). The level of activation then depends on the expression strength of the upstream gene (Fig. 3C); the experimentally observed value is obtained for a moderate rate of one transcription event every 5 min. Altogether, these results show that our modeling quantitatively reproduces the effect of TSC on well-defined constructions on the chromosome; we can now turn our attention to its contribution to the global regulation by SC along entire genomes.

### 3.3 Topoisomerase distribution and regulatory activity on the chromosome is dictated by genomic architecture

As shown above, adjacent genes interact on the chromosome through TSC, according to complex yet predictable rules that already emerge from our simplified modeling. Note that this observation goes against the classical dogma of gene regulation centred on transcription factors, where a gene’s expression is entirely determined by its promoter sequence and independent from, *e.g.*, its location on the genome and neighbouring genes [40]. Considering the density of bacterial genomes, we may thus expect that the mechanical constraints induced by transcription of adjacent genes, and tightly related to their orientation, play an important role in the genome-wide coordination of gene expression. To directly test this effect, the local distribution of supercoils along the chromosome should be measured, but the only available data of this kind lack sufficient spatial resolution [41]. However, an indirect measurement was provided by ChIP-Seq for the two major topoisomerases in *Mycobacterium tuberculosis* [39]. The analysed distributions indicate that topoisomerase I and the DNA gyrase bind at a higher rate in divergent and convergent regions respectively (Fig. 4A). Although these distributions might be affected by factors not considered in our modeling (gyrase high-affinity sequences, 3D conformation of DNA, ChIP-Seq protocol), these observations are in full agreement with our expectations based on a simplified description of TSC, *i.e.*, the two topoisomerases are more active in convergent/divergent regions because of the different SC levels induced by neighbouring transcription rather than because of any sequence specificity (Fig. 1). In order to simulate and reproduce such *in vivo* genome-wide data (Fig. 4B), we reasoned that simulating an entire chromosome would involve many arbitrary choices (definition of topological domains and transcription units, gene expression strengths…). Since most observed properties depend primarily on gene orientations, more than on precise gene positions and distances, we rather decided to simulate a 30-kb-long toy model circular genome involving three topological domains (Fig. 4C), each of them carrying two identical 1-kb-long active genes (yellow) in different configurations, and a central inactive gene (grey) which does not experience any transcription elongation but is used as a regulatory probe. All promoters have the same basal initiation rate and SCdependence: any observed difference in the initiation rate of inactive promoters thus directly reflects the effect of TSC induced by their neighbours, according to their orientation.

**Figure 4:**
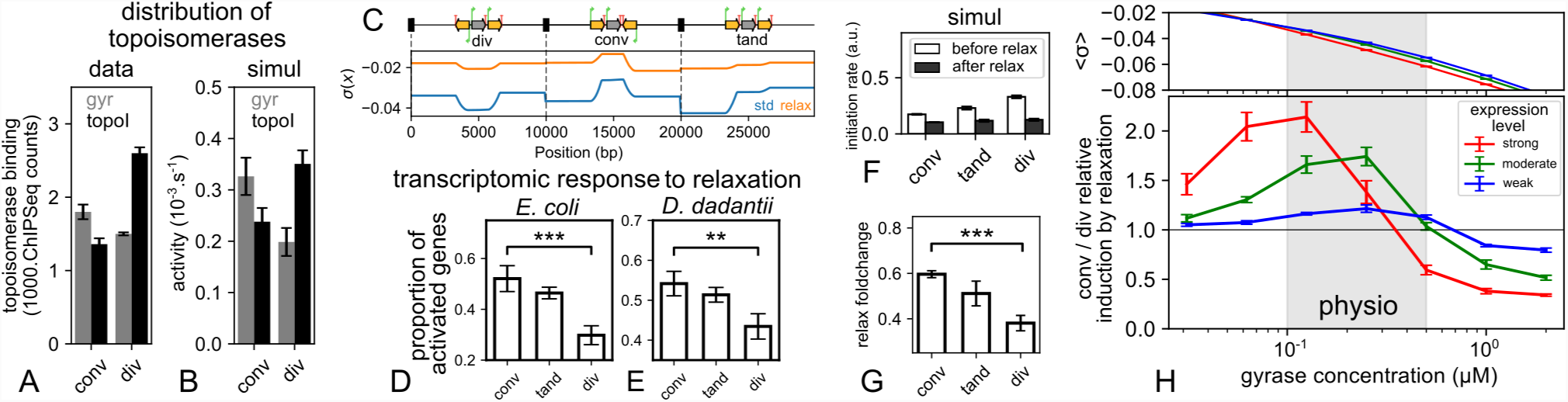
**(A)** ChIP-Seq data show that topoisomerase I and the DNA gyrase preferentially bind in divergent and convergent regions, respectively [39]. **(B)** The observed distribution naturally emerges from our modeling of TSC in physiologically relevant conditions (moderate gene expression and 0.25 *µ*M gyrase concentration, see Materials and Methods). **(C)** Simulations were carried on a model genome with three distinct topological domains, each carrying two active genes (yellow) in different orientations, and a central inactive gene (grey) as a regulatory probe. The heterogeneous SC levels are shown before (blue) and after (orange) relaxation by gyrase inhibition, and they give rise to the heterogeneous recruitment of topoisomerases observed in B. **(D)** Transcriptomics data in *Escherichia coli* from [8] show that genes’ response to chromosomal relaxation is tightly related to their local orientation. **(E)** The same is observed in new RNA-Seq data from *Dickeya dadantii*, in exponential as well as stationary (Fig. S3B) phase, confirming that this feature is not organismor condition-specific. **(F)** Simulating the action of the antibiotics exhibits a different effect on the promoters’ initiation rates, depending on their local orientation. **(G)** As a result, relaxation favours convergent genes versus divergent ones, in agreement with genome-wide experimental data. **(H)** The average SC level (upper panel) and relative convergent/divergent foldchange due to relaxation (lower panel) was computed for a range of initial DNA gyrase concentrations and expression strengths (of all active genes), corresponding to effective waiting times of 1, 2.5 and 10 min between transcription events at each promoter, respectively. Data shown in B, C, F and G correspond to the central datapoint with moderate expression.

The relatively small differences in topoisomerases binding observed between convergent and divergent regions (Fig. 4A) are reproduced (Fig. 4B) when the genes are moderately expressed in these simulations (one transcript every 2.5 minutes), as expected for most of them in the genome. In bacteria, the regulatory effect of SC is generally analysed from transcriptomes obtained shortly after chromosome relaxation by antibiotics (novobiocin, norfloxacin) which inhibit the DNA gyrase [7, 8, 9, 10]. Based on the previous observations, the picture of a global relaxation might yet appear as slightly misleading, if the effect of gyrase inhibition is very different in convergent vs divergent regions. In this case, spatial topological heterogeneities resulting from TSC may play an important role in the differential response of genes to a global change in gyrase activity, which is itself a key actor in the cellular response to environmental variations [2, 5]. To test this hypothesis, we analysed several transcriptomes obtained under different conditions of chromosomal relaxation shock. In *E. coli*, they were obtained by microarrays, either from mutant strains favouring a strong relaxation by norfloxacin [8] (Fig. 4D), or with novobiocin in the wildtype strain [7] (Fig. S3A in Supp. Mat.). In both cases, a significant difference is observed in the response of convergent vs divergent genes (*P* = 2.3 × 10^−4^ and *P* = 0.02 respectively, *χ*^2^ test). Since we are investigating a mechanism relying on basal properties of transcription shared by many different bacterial species, we assessed the universality of this observation by conducting RNA-Seq experiments on *Dickeya dadantii*. Transcriptomes were collected 15 minutes after a novobiocin shock applied either in exponential phase (Fig. 4E) or, for the first time in enterobacteria, at the transition to stationary phase (Fig. S3B), and the same difference is observed as in *E. coli* (*P* = 8 × 10^−3^ and *P* = 1.6 × 10^−3^, respectively). Altogether, in spite of strong differences in organism, experimental conditions and methods, a systematic and coherent relation between gene orientation and transcriptional response is observed in all data. But interestingly, while we intuitively expected convergent genes to be more hampered by gyrase inhibition (since the latter is more active in these regions), the reverse effect is observed, as they systematically appear more activated by chromosome relaxation than divergent genes. This effect was already noted by Sobetzko [13], who invoked a putative difference in promoter sequences, suggesting that evolution has placed relaxation-activated promoters in convergent regions where DNA is more relaxed. In view of our previous observations, we rather propose that this behaviour could emerge even if most promoters exhibit exactly the same “standard” SC-sensitivity, as a (counter-intuitive) result of the redistribution of SC resulting from gyrase inhibition. Because of the complexity of the process, this scenario cannot be easily predicted, but it can be tested within our modeling where all promoters have the same response curve (Fig. 1C) and gyrase inhibition can be simulated explicitly. Using the same parameters as in Fig. 4B, we simulate one hour of transcription, and then mimic the action of antibiotics by suddenly dividing the gyrase concentration by 5; the initiation rate before and after the shock are then compared (Fig. 4F). As expected, all initiation rates are reduced by the global relaxation of the chromosome, but by different factors: more than 2.5 for divergent genes, against less than 2 for convergent genes (Fig. 4G). Remarkably, the shock thus leads to a redistribution of RNAPs in (relative) favour of convergent genes; since gene expression levels are normalised in a transcriptomics experiment, the simulations thus predict a stronger activation of the latter in such data, precisely as systematically observed. Interestingly also, the simulation makes it possible to analyse the mechanism responsible for this counterintuitive behaviour, *i.e.*, the different SC levels in the convergent vs divergent regions before and after the shock (Fig. 4C). Before the shock, the convergent gene was already located in a relaxed region because of TSC, and was thus already partly repressed, whereas the divergent gene was in a strongly negative region and thus fully active (blue curve). In contrast, after the relaxation shock, both of them have been shifted to a similar repressed state; the amount of SC relaxation (vertical distance between the blue and orange curves) is much stronger for divergent genes, and also its repressive effect.

### 3.4 Interplay between gene expression strength and topoisomerase activity in global regulation by TSC

Since generic rules of basal regulation by TSC seem well-reproduced by our model, we can now use it to explore how these rules and the whole system behaviour depend on the precise conditions of the simulation. We tested the influence of two biologically relevant parameters: (1) gene expression strength (of all active genes in our model genome), and (2) DNA gyrase concentration. The latter parameter actually also mimics the controlled variations of gyrase activity in the cell in response to environmental constraints, in particular through variations of the ATP/ADP ratio in exponentially growing vs stationary cells [24]. Fig. 4H shows the average superhelical density and the relative convergent/divergent activation factor due to gyrase inhibition, for different gyrase concentrations and basal expression rates, corresponding to effective waiting times of around 1, 2.5 and 10 min between transcription events of each active gene respectively. Unsurprisingly, the negative SC level (upper panel) increases with gyrase concentration; but interestingly, this level also strongly depends on the gene expression strength, with differences as large as Δ*σ* ≃ 0.01 for physiological levels of DNA gyrase. This difference is of the same order as the measured effects of novobiocin [3] or topoisomerase mutations (see below). Actually, such a behaviour has already been observed experimentally on a plasmid, which was found more supercoiled in the cell when it carried a strong than a weak promoter [42]. Our simulations show that it results from the nonlinear response curve of topoisomerases in presence of TSC (Fig. 1D): since highly expressed domains experience strong transient SC inhomogeneities, topoisomerases sometimes bind very actively and non-uniformly in these domains, resulting in a stationary stronger SC level. Significant consequences result from this observation: (1) in a living cell, the local SC level is likely higher in strongly than weakly expressed topological domains (in addition to orientation-dependent effects already noted); (2) the SC level measured on a plasmid (*e.g*., in a chloroquine gel experiment) only semi-quantitatively reflects the chromosomal level, as both levels are affected by the presence/absence and activity of promoters.

Altogether, these observations show that defining the “supercoiling-sensitivity” of promoters purely from transcriptomics experiments is delicate, since the SC levels *locally* experienced by these promoters on the chromosome may differ quite strongly from the *global* levels measured from plasmids, and even more so when considering other factors disregarded here, such as nucleoid-associated proteins [2, 5, 36]. Note that the SC levels observed in the simulations are somewhat lower than those usually measured *in vivo* on plasmids, which is consistent with our neglecting the writhe contribution (and keeping in mind the previous reservations). Simulations of gyrase inhibition (Fig. 4H, lower panel) also show that the relative induction of convergent vs divergent genes (datapoints above 1) is observed at all gene expression levels (and more strongly for highly expressed genes), but not when the initial gyrase concentration is very high. In particular, strongly expressed convergent genes might even be repressed by gyrase inhibition in physiological conditions (right part of red curve), as one would expect if these genes specifically require DNA gyrase to stay active. This effect would probably be even stronger had we considered the stalling effect of positive supercoils on transcription elongation [17].

### 3.5 TSC-mediated regulation in experimental evolution

If TSC constitutes an ancestral mode of regulation, it might play an important role in genome evolution. An interesting example is that of synteny segments, *i.e.*, genes remaining adjacent in evolution, which exhibit distinct orientation and co-expression features currently unexplained [43]. Here, we focus on the longest-running evolution experiment, during which SC was indeed identified as a critical adaptive actor [6]. In this experiment, 12 independent *E. coli* populations were grown by serial daily transfer in minimal medium for 50,000 generations [28]. In one representative population, fitness already strongly increased after 2000 generations (Fig. 5A) together with the SC level within the cells (by Δ*σ* ≃ −0.008 as measured on a plasmid, Fig. 5B). These changes result from only 6 mutations [30], including one beneficial SNP in the *topA* gene encoding topoisomerase I, which reduces its efficiency. A second increase in SC (of an additional Δ*σ* ≃ −0.003) occurred before 20,000 generations, owing to the fixation of another beneficial mutation affecting *fis* among 45 mutations in total [6]. It was proposed that SC-modifying mutations provided an efficient way to change the cell expression programme globally [6]. We then expect TSC to contribute in this transcriptional response, probably in an opposite direction to that observed in relaxation shocks (since the SC level is here increased).

**Figure 5:**
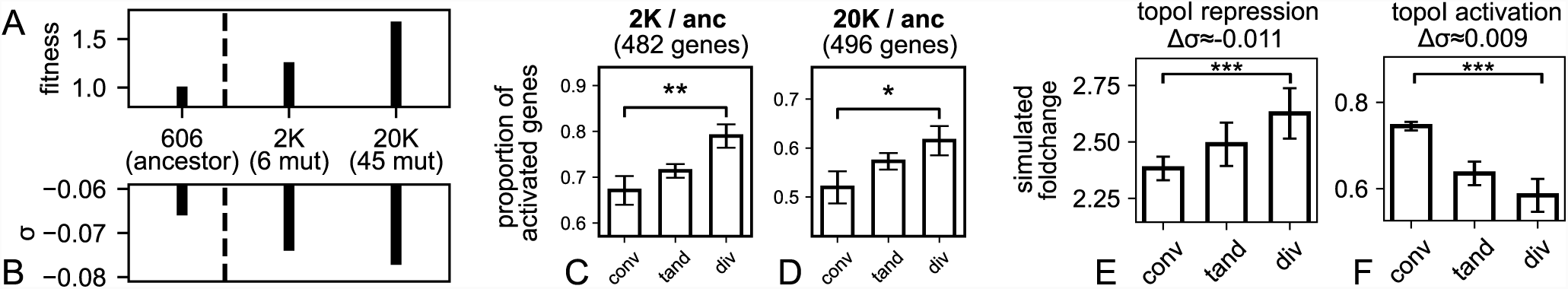
Gene expression profiles in clones from the long-term evolution experiment with *E. coli* [27, 28] exhibits the signature of TSC-mediated regulation. **(A)** 2K and 20K clones from the Ara-1 population evolved increased fitness relative to their ancestor after 2000 and 20,000 generations, respectively (data from [6]). **(B)** The SC level sequentially increased, owing to the selection of two SC-modifying mutations (data from [6]), first in *topA* (among 6 mutations in 2K), then in *fis* (among 45 mutations in 20K, including the *topA* mutation) [30]. **(C)** In 2K, divergent genes are more activated than convergent ones with respect to the ancestor (*P* = 1.6 × 10^−3^), as predicted by TSC. Differentially expressed genes were selected with a loose threshold (see text): we indicate the number of convergent+divergent responding genes. **(D)** Same for the 20K clone (*P* = 0.016). **(E)** We simulated the mutation in *topA* by reducing the activity of topoisomerase I by 2-fold: as expected and observed *in vivo*, divergent genes are favoured by the resulting increase in negative SC. **(F)** Conversely, a 2-fold increase in topoisomerase I activity favours convergent genes, in the same manner as gyrase inhibition (Fig. 4G).

To test this putative effect of TSC, we collected the global transcription profiles of the ancestral and evolved strains using microarrays (see Materials and Methods). Evolved strains exhibit differentially expressed genes with respect to the ancestor, yet their number is relatively low under standard statistical selection procedures compared to relaxation shock experiments (97 and 148 genes in the 2K and 20K strains, respectively; q-value *<* 0.1). However, in contrast to the latter case, here the alterations in gene expression are not caused by a fast change in SC, but rather by an inheritable SC change, the consequences of which can be balanced by other actors of the regulatory network, including transcription factors. Since the few identified genes with altered expression are those where the activation/repression is strongest, most of these changes are probably due to the strong and localised action of “digital” (on/off) transcription factors. Indeed, SC alone usually exhibits a more subtle and “analogue” (more/less) regulatory effect on a larger number of genes [2, 40]. To analyse the latter effect, we therefore generated new datasets of differentially expressed genes, in numbers similar to those obtained in relaxation shock experiments, by reducing the threshold of statistical significance (Fig. 4, p-value *<* 0.25). This operation implies that these larger datasets likely contain many false positives which might blur out the investigated effects of TSC, but the latter may still emerge as a statistical feature. We therefore looked for a statistical relationship between gene orientation and response (Fig. 5 C-D). Both evolved clones with increased SC level indeed exhibit a higher rate of activated divergent genes, as expected from our previous analysis. The confidence level is high for the 2K strain where there are few other mutations (*P* = 1.6 × 10^−3^), and also significant for the 20K strain (*P* = 0.016) which exhibits more mutations that could contribute to rewire the regulatory network independently of SC effects. Note that the observed behaviour does not depend on the precise significance threshold chosen above and is robust with more stringent selection of differentially expressed genes, although with weaker statistical significance owing to their lower number (data not shown). To crosscheck if it is indeed reproduced by the model, we ran simulations involving decreased (Fig. 5E) or increased (Fig. 5F) topoisomerase I activity (by 2-fold, compatible with the measured SC variation Δ*σ* ≃ −0.011). They confirmed a relationship between relative activities of topoisomerases and orientation-dependent response of genes. Altogether, these analyses show that (1) TSC defines robust rules that allow the cell to selectively activate convergent or divergent genes by tuning its global SC level, independently from more specific regulation of each promoter; (2) experimentally evolved populations in which SC-modifying mutations were selected exhibit changes in global transcription profiles consistent with the TSC mode of regulation; (3) at larger evolutionary timescales, TSC might also contribute to the selection of events such as insertions/deletions or inversions, which could modify the expression profile by changing the relative distances and orientations between adjacent genes, even without modifying the gene coding or promoter sequences.

## 4 CONCLUSION

Many bacterial genes exhibit “supercoiling sensitivity” in transcriptomics experiments, the mechanisms of which remain unclear. Inspired by classical regulation models, most efforts have focused on promoter sequences to explain their response [7, 2]. Here, we propose alternatively that strong spatial variations of DNA supercoiling along the chromosome equally contribute to the complexity of this response, and allow the cell to activate its genes in a global but selective way depending on their local orientation. Such variations were measured at low resolution in the bacterial chromosome [41], and were already linked to gene orientations in eukaryotes [18], but these effects remained nonetheless as yet neglected in quantitative regulatory models. Those data obtained in very different organisms suggested that the fundamental mechanical properties of transcription give rise to an ancestral coupling between genome expression and torsion (here called TSC). This coupling underpins a basal form of regulation that might affect all organisms, albeit with different rules owing to differences in genome structures, topoisomerase enzymes, etc. In bacteria, we have analysed here some of these complex yet predictable rules by which global variations of DNA topology are distributed at gene promoters, and allow for a finetuned global regulation mode under the control of cell physiology. The model presented, which was kept voluntarily as simple as possible, is already remarkably predictive of how the genome architecture dictates this regulation by mechanically coupling adjacent genes because of the transcription process itself, and depending on their relative orientations. As a further step, the model may be used to directly simulate a larger chromosomal region *in vivo*, which will raise new substantial issues regarding the proper definition of topological barriers, transcription units, and other required components. An especially attractive application is the so-called “pathogenicity islands” where virulence genes are co-localised and remain co-regulated when they are horizontally transferred from one pathogenic species to another. The model predictions should then be experimentally tested with more detail and in a wider range of bacterial organisms. Such analyses will likely exhibit more subtle regulatory effects involving nucleoid-associated proteins and other actors disregarded here, which contribute in the 3D organisation of the chromosome [36]. We anticipate that this comparison will lead to an incremental refinement of the modeling (in particular by incorporating sequence-dependent properties of DNA and DNA-protein interactions), toward a more comprehensive description of the global transcriptional regulation embedded in the bacterial chromatin structure.

## Supplementary data

Supplementary data are available online.

