## Supplementary Material for "Bacterial genome architecture shapes global transcriptional regulation by DNA supercoiling"

Bilal El Houdaigui, Raphaël Forquet, Thomas Hindré, Dominique Schneider,
William Nasser, Sylvie Reverchon and Sam Meyer

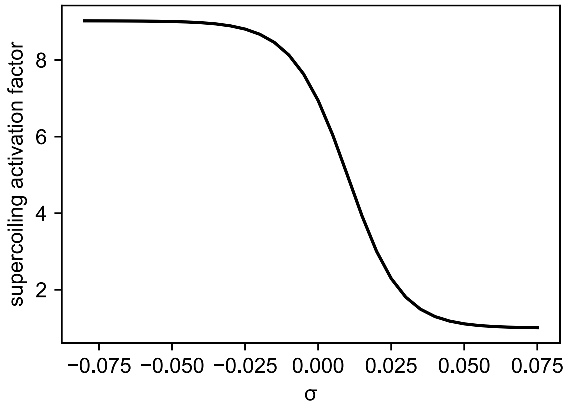


Figure 1: Shifted effective promoter activation curve used to simulate the data of Chong et *al.* [1] (Fig. 2B). This modified curve accounts for the stalling effect of positive supercoils on transcription elongation in this *in vitro* experiment in absence of DNA gyrase, and is consistent with the observed repressive effect of positive supercoils.

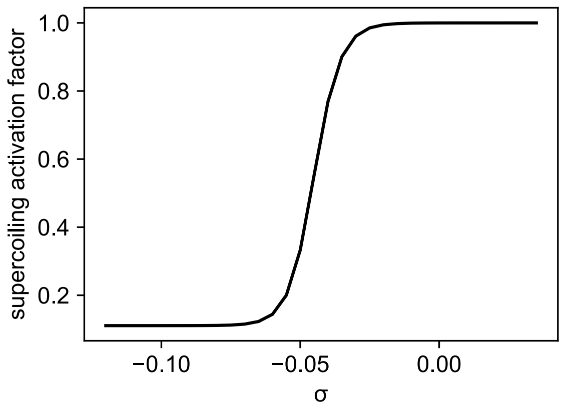


Figure 2: Reversed promoter activation curve used for the *gyrA* promoter (Fig. 3B), which is a very specific promoter activated by DNA relaxation to ensure an homeostasis of the SC level in the cell [2].

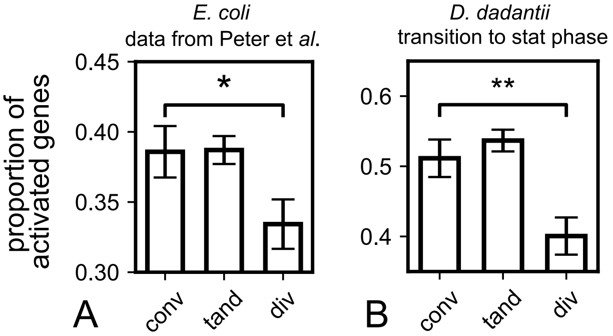


Figure 3: Additional analyses of genes' response to DNA relaxation. **(A)** Same as Fig. 4D, for the novobiocin data of Peter et *al.* [3]. For this experiment, the authors only provided global p-values obtained from many experimental conditions involving topological variations, resulting in a low number of significantly affected genes. For consistency with the other data presented here, we focused on the response to novobiocin treatment, for which the gene expression levels were provided but not the activation p-values. We therefore considered all genes as activated or repressed, explaining that the proportions are much closer to each other in this graph (vertical scale), but with much narrower confidence intervals (larger number of genes in the dataset). **(B)** Same as Fig. 4E, for the transcriptome obtained at the transition to stationary phase. Although the global expression pattern is very different in this phase, the effect of relaxation is similar for convergent vs divergent genes.
